# Stridulations of *Melolontha spp.* larvae open up new possibilities for species-specific pest monitoring in soils

**DOI:** 10.1101/520734

**Authors:** Carolyn-Monika Görres, David Chesmore

## Abstract

Root-feeding Scarabaeidae larvae can pose a serious threat to agricultural and forest ecosystems, but many details of larval ecology are still unknown. We developed an acoustic data analysis method for gaining new insights into larval ecology. In a laboratory study, third instar larvae of *Melolontha melolontha* (*n*=38) and *M. hippocastani* (*n*=15) kept in soil-filled containers were acoustically monitored for 5 min each, resulting in the first known stridulation recordings for each species. Subsequent continuous monitoring of three *M. hippocastani* larvae over several hours showed that a single larva could stridulate more than 70 times per hour, and stridulation rates increased drastically with increasing larval abundance. The new fractal dimension-based data analysis method automatically detected audio sections with stridulations and provided a semi-quantitative estimate of stridulation activity. It is the first data analysis method specifically targeting Scarabaeidae larvae stridulations in soils, enabling for the first time non-invasive species-specific pest monitoring.

**Key message:**

- Root-feeding scarab beetle larvae, known as white grubs, can be serious agricultural and forest pests.
- White grub infestation monitoring is difficult due to their cryptic lifestyle, but detailed monitoring data is essential for effective pest control.
- We present the first acoustic data analysis routine targeting stridulations, using *Melolontha melolontha* and *M. hippocastani* as model organisms.
- This new method provides for the first time the basis for the development of tools for non-invasive, species-specific, and rapid white grub monitoring in soils.

## Introduction

The Melolonthinae, a subfamily of the scarab beetles (Scarabaeidae), comprises 29 tribes (Smith 2006) of which several pose a serious threat to agricultural and forest ecosystems all over the world (e.g. Keller and Zimmer 2005; Jackson & Klein 2006; Frew et al. 2016). The adult beetles have strong mandibles for mainly eating leaves, however serious damage to trees and agricultural crops is first and foremost caused by the soil-dwelling, root-feeding larvae, also known as white grubs (Jackson and Klein 2006). In Europe, two species of Melolonthinae which have to be considered as important pest insects are *Melolontha melolontha* (Common Cockchafer) and *M. hippocastani* (Forest Cockchafer) (Keller and Zimmer 2005). These two species are very similar in their biology and are not host plant specific. While *M. melolontha* can be found in more open habitats (e.g. pastures, vegetable crops, orchards, vineyards), *M. hippocastani* mainly thrives in deciduous forests (Wagenhoff et al. 2014; Sukovata et al. 2015). Currently, these two species occur as pests in Austria, Czech Republic, France, Germany, Italy, Poland and Switzerland, but the available monitoring data is incomplete (Keller and Zimmer 2005), and *Melolontha spp.* outbreaks are expected to spread again after a decline in population sizes in the middle of the last century (Kahrer et al. 2011). The reasons behind the recovery of *Melolontha spp.* populations are mainly unknown (Kahrer et al. 2011), but are of serious concern as e.g. infested forest areas become more susceptible to droughts and secondary diseases, and forest regeneration can be hindered (Immler and Bussler 2008; Wagenhoff et al. 2014; Sukovata et al. 2015).

Strategies to control white grub infestations of *Melolontha* spp. include mechanical (soil-covering nets, ploughing), chemical (pesticides) and biological (*Beauveria* spp., nematodes) measures. These have been applied to various degrees of success during the past century, but until today, there is no generally applicable, environmentally friendly pest control strategy to control white grub infestations at agricultural and forest sites (Jackson and Klein 2006; Woreta 2015). One of the main reasons for this are the difficulties associated with monitoring these soil-dwelling insects (Johnson et al. 2007). Due to their cryptic lifestyle, white grubs can live for years unnoticed in the soil. The larvae of *M. melolontha* and *M. hippocastani* live three and four years, respectively, in the soil until they develop into the adult chafer. During this time, the larvae pass three different instars. They only reach pest status if the larval abundance surpasses a certain threshold (Immler and Bussler 2008; Frew et al. 2016).

The current standard method for monitoring larval abundances and to confirm white grub infestations is to excavate the soil. This type of monitoring is invasive, laborious and time-consuming and cannot be performed at a high temporal frequency (Johnson et al. 2007; Immler and Bussler 2008; Mankin et al. 2009). As a result, *Melolontha* spp. infestations can go unnoticed until severe damage to the vegetation becomes visible, larval abundances can be underestimated, and many details of the larval ecology are still unknown. However, a stringent monitoring and a detailed knowledge of a target species’ ecology are essential for the development of environmentally friendly and successful pest control measures (Frew et al. 2016).

One way to improve the monitoring of *Melolontha* spp. larvae is the use of acoustics. Acoustic sensors inserted into the soil have already been successfully applied for detecting white grub infestations and also for the quantification of larval abundances non-invasively (Zhang et al. 2003; Mankin et al. 2009a; Mankin et al. 2011). However, these studies were purely based on monitoring incidental sounds (movement, feeding sounds) and species identification still had to be confirmed via soil excavations. Several different species belonging to the Melolonthinae can co-occur in soils – not all of them necessarily being regarded as pest species (Jackson and Klein 2006) – and even for experts it can be difficult to differentiate the species rapidly in the field when they are still in their larval stage. Especially differentiation of *M. melolontha* and *M. hippocastani* larvae based only on morphological features alone does not seem possible (Krell 2004). One promising way to overcome this problem is to shift the focus of acoustic monitoring from incidental sounds to stridulations. Stridulations are actively produced sounds for communication created by rubbing together certain body parts (Alexander and Moore 1963). Harvey et al. (2011) have shown for three different Scarabaeoidea species that the larvae produce species-specific stridulation patterns. Targeting larval stridulations has the potential to greatly improve pest monitoring by enabling non-invasive, species-specific monitoring with high spatial and temporal coverage. However thus far, stridulations have rarely been studied and never been utilized in field monitoring programs (Sprecher-Uebersax & Durrer 1998; Harvey et al. 2011). This study presents the first stridulation audio recordings of *M. melolontha* and *M. hippocastani* larvae and demonstrates their applicability for easy identification of these two species. We also designed a new data analysis routine for rapid detection and quantitative estimation of *Melolontha* spp. stridulation events in soil audio recordings based on fractal dimension. *Melolontha* spp. were chosen due to their pest status in Europe, but they also serve as model organisms for white grubs in general. The acoustic data analysis method presented here should be easily transferable to other soil-dwelling Melolonthinae and Scarabaeidae species, providing for the first time the basis for the development of tools for non-invasive, species-specific, and rapid pest monitoring in soils, but also for soil biodiversity and soil insect ecology studies in general.

## Material and methods

### Cockchafer larvae for acoustic laboratory measurements

Acoustic measurements were performed with scarab beetle larvae of the species *Melolontha hippocastani* (Forest cockchafer) and *M. melolontha* (Common cockchafer). Thirty *M. hippocastani* larvae (second instar) were excavated in a mixed coniferous forest on sandy soil (Hessisches Ried, Pfungstadt, Germany) in November 2015, and individually kept in small plastic containers (100 ml) with perforated lids in the laboratory in the dark at near constant room temperature (∽17 °C). Each container was filled with soil from the excavation site. Approximately every two weeks, carrot slices were added to the containers as food source and the soil sprayed with tab water to keep it from drying out. In the laboratory, the larvae shed their exoskeleton once passing from second to third instar in March 2016. In June 2016, 15 larvae were randomly selected for acoustic monitoring in the laboratory. Seventy-five third instar larvae of *M. melolontha* were excavated in a meadow on sandy soil (Blaubeuren-Weiler, Germany) in May 2017, and transferred to the laboratory being kept in the same way as the *M. hippocastani* larvae. One week after excavation, 38 of these *M. melolontha* larvae were randomly selected for acoustic monitoring.

### Acoustic monitoring

All acoustic measurements were conducted at room temperature (∽20 °C) in a plastic box (60 cm x 40 cm x 33 cm) insulated with acoustic foam to reduce background noises. Detailed instructions for replicating the acoustic sensors described in the following paragraphs can be obtained from David Chesmore. In the first experiment, each larva was acoustically monitored for 5 min by placing one plastic container at a time into the box with a low-cost sensor attached to the outside wall of the container. The sensor was self-made based on a piezoelectric transducer (amplified, gain 20). It was connected to an external battery box which in turn was plugged into the microphone input of a commercially available audio recorder (TASCAM Linear PCM Recorder DR-05 Version 2, TEAC Europe GmbH, Wiesbaden, Germany). Sounds were recorded in .wav format with an audio sampling rate of 44.1 kHz. Audio recordings of *M. hippocastani* and *M. melolontha* were conducted in June 2016 and June 2017, respectively. For the *M. hippocastani* recordings, the box was placed in a room with almost no background noises, whereas for the *M. melolontha* recordings, the box was placed in a laboratory with a noisy environment.

After the acoustic screening of the *M. hippocastani* larvae, the three most sound-producing larvae from this population were selected for a second experiment. An acoustic sensor consisting of a piezoelectric transducer encased in a water-proof, silicone sealed plastic case (length: 21 cm, width: 3 cm, thickness: 0.5 cm) was positioned upright in a glass jar (volume: 2.7 l, height: 24 cm, diameter: 10 cm). Subsequently, the glass jar was completely filled with sandy soil from the original excavation site of the larvae. In the first step of the experiment, one of the three selected *M. hippocastani* larvae was placed on top of the soil together with fresh carrot slices as food source. The larva had to burrow itself into the soil. Upon placing the larva in the glass jar, sounds were recorded continuously with the buried sensor for 12 hours. A new audio file was created every 50 min. Two weeks later, the remaining two selected *M. hippocastani* larvae were also added to the jar together with fresh carrot slices. A new continuous audio recording was started, this time lasting for 18 hours. Apart from the sensor, the audio recording equipment and the audio sampling rate were the same as in the first experiment. During the recordings, the silent box was stored in a room with little background noise.

### Data analysis

In a first step, all audio files were bandpass filtered (200 – 5000 Hz) and normalized (maximum amplitude −1.0 dB, DC offset removed) using the software Audacity 2.1.3 (Audacity Team 2017). Afterwards, stridulation events were manually detected and counted by visual and audible inspection of each audio file. In a second step, a new data analysis routine utilising fractal dimension was developed for automated rapid detection and quantitative estimation of stridulation events based on the work of Schofield (2011). The data analysis routine was written with the software R 3.4.3 (R Core Team 2017) utilizing the R packages “fractaldim_0.8-4” (Sevcikova et al. 2014) and “tuneR_1.3.2” (Ligges et al. 2016) (see Online Resource 1). It was applied to all audio files generated in the second experiment after these files were pre-processed in Audacity as described.

In fractal dimension analysis, the waveform of an audio recording is considered as a geometric shape and its complexity approximated through the calculation of scalar values (Schofield and Chesmore 2008). The geometric complexity of stridulation events differs from incidental larval sounds (movement, feeding sounds) or background noise. First, the pre-processed audio files were imported into R and sliced into 2 s long sections (*sc*), of which only the amplitude values were extracted for further analysis (Fig. 1a). For each *sc*, fractal dimensions (*D*) for the waveform were calculated twice with the madogram estimator (Gneiting et al. 2010) and non-overlapping frames (*f*) using a frame size of 88.2 samples (= 2 ms) and 176.4 samples (= 4 ms), respectively. Subsequently for each *f* in *sc, D* was converted into a fractal distance (*FD*) by calculating its median deviation from the median (md) (Leys et al. 2013):

**Fig. 1.**
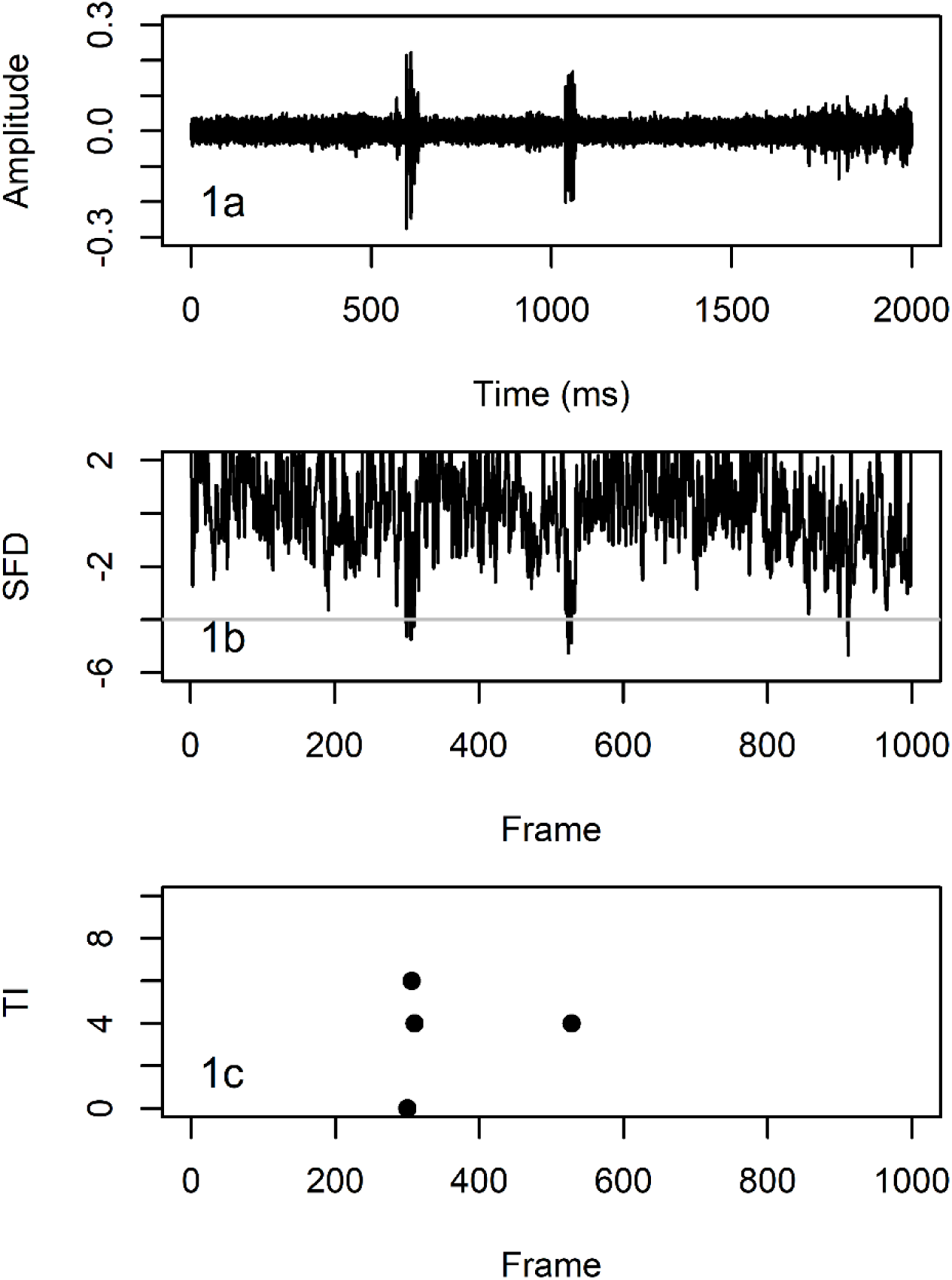
Detection of cockchafer larvae stridulations using fractal dimension analysis (see text for details). Fig. 1a: Audio recording with two stridulations (at ∽650 ms and ∽1100 ms) and larval moving sounds (from ∽1700 ms onwards). Fig. 1b: Summed fractal distance (*SFD*) for every 2 ms (=frame) of the audio recording. Peaks crossing a threshold of −4.0 are first indicators of stridulation events. Fig. 1c: Number of frames between adjacent peaks (*TI*) crossing the threshold in Fig. 1b. A distance of less than 10 frames is indicative of a peak of clusters crossing the threshold in Fig. 1b, and thus a stridulation event

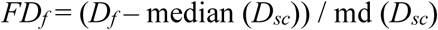

The *FD* timeseries with a frame size of 88.2 samples consisted of 1000 samples for each *sc*. The *FD* timeseries with a frame size of 176.4 samples was linearly interpolated to comprise the same amount of samples, and then both timeseries were summed up (*SFD*) (Fig. 1b). The *SFD* timeseries was utilized for the detection and quantitative estimation of stridulations. First, the R function ‘rle’ (= run length encoding) was used to filter out all *f* with *SFD* > −4.0. Second, all *f* were filtered out where *SFD* < −4.0 for more than 2 adjacent *f*. Third, the time interval (in *f*) between the remaining *f* was calculated (*TI*). Clusters of single *f* with *SFD* < −4.0 being spaced apart less than 10 *TI* within the clusters were indicative of stridulation events (Fig. 1c). For each 50 min audio file, stridulation activity (*STRAC*) was automatically estimated by multiplying *TI* 1 to 10 with their respective frequencies and summing up the products.

## Results

### Stridulation patterns

Stridulations by the two species were easily recognizable when listening to the audio recordings. The common stridulations of *M. melolontha* and *M. hippocastani* consisted of short bursts of sound (Fig. 2). They were very similar and both peaked at a frequency of ∽1700 Hz, however, *M. melolontha* stridulations were of longer duration than the ones of *M. hippocastani*. Stridulations often occurred in pairs, but repetitions up to 4 times were also recorded. A second type of stridulation which was recorded less frequently from both species consisted usually of 4 (seldom only 2 or 3) repeated patterns with a duration of ∽250 ms each and a frequency peak at 3000 Hz (Fig. 3).

**Fig. 2.**
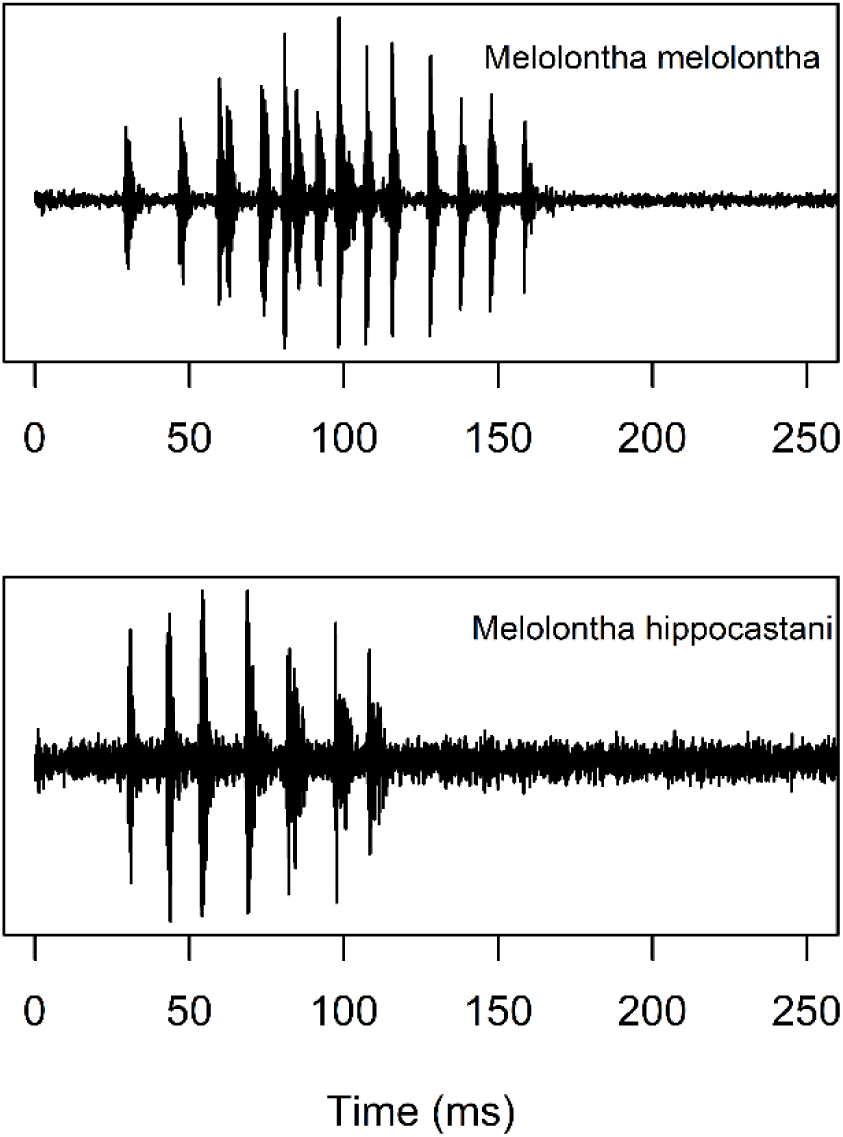
Comparison of acoustic patterns produced by stridulation of larvae (third instar) of *Melolontha melolontha* and *M. hippocastani*

**Fig. 3.**
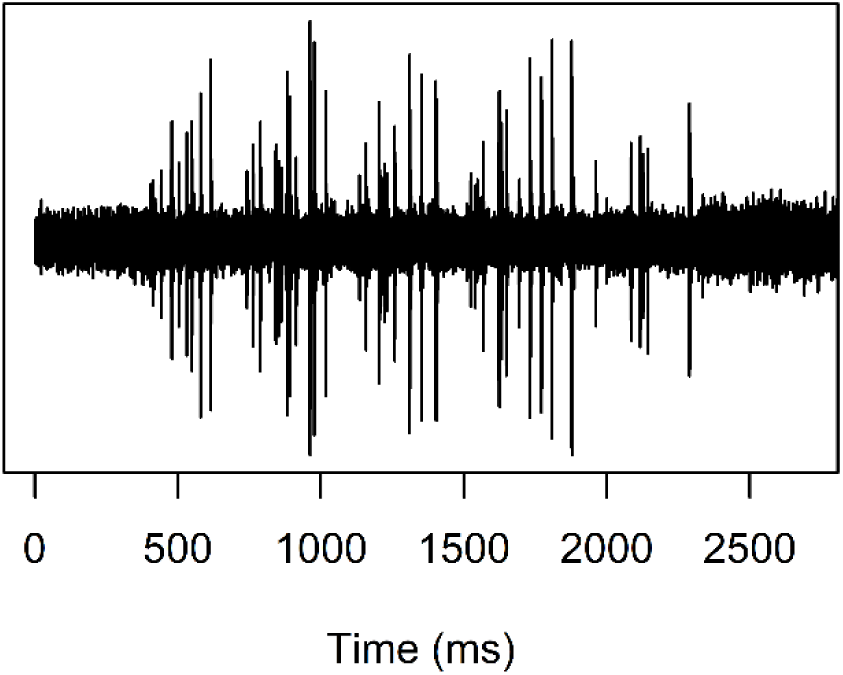
Acoustic pattern produced by stridulation of a third instar *Melolontha hippocastani*

### Distribution of stridulation events

During the 5 min acoustic screening of individual larvae, only few stridulations were detected. In total, only 5 stridulations from 3 different individuals were recorded from the set of 15 *M. hippocastani* larvae. Of the 38 *M. melolontha* larvae, 5 individuals were caught stridulating, producing 16 stridulations altogether. In contrast, numerous stridulations were observed during the continuous acoustic monitoring of *M. hippocastani* larvae in the second experiment (Fig. 4). The first larva which was placed in the soil-filled glass jar stridulated 75 times in the first 50 min after placement. The stridulation rate dropped to 15 stridulations h^-1^ over the next 4 h and subsequently, the larva almost completely stopped stridulating except for a few single stridulation events. In total, the larva produced 188 stridulations during 12 h of continuous recording. Stridulation activity drastically increased in the soil-filled glass jar after adding 2 more larvae. In the first 2.5 h alone, the 3 *M. hippocastani* larvae stridulated 682 times. Afterwards, the stridulation rate levelled below 70 stridulations h^-1^ with periods of high activity alternating with periods of very low activity. Over the course of 18 h of continuous recording, the 3 larvae produced 1100 stridulations.

**Fig. 4.**
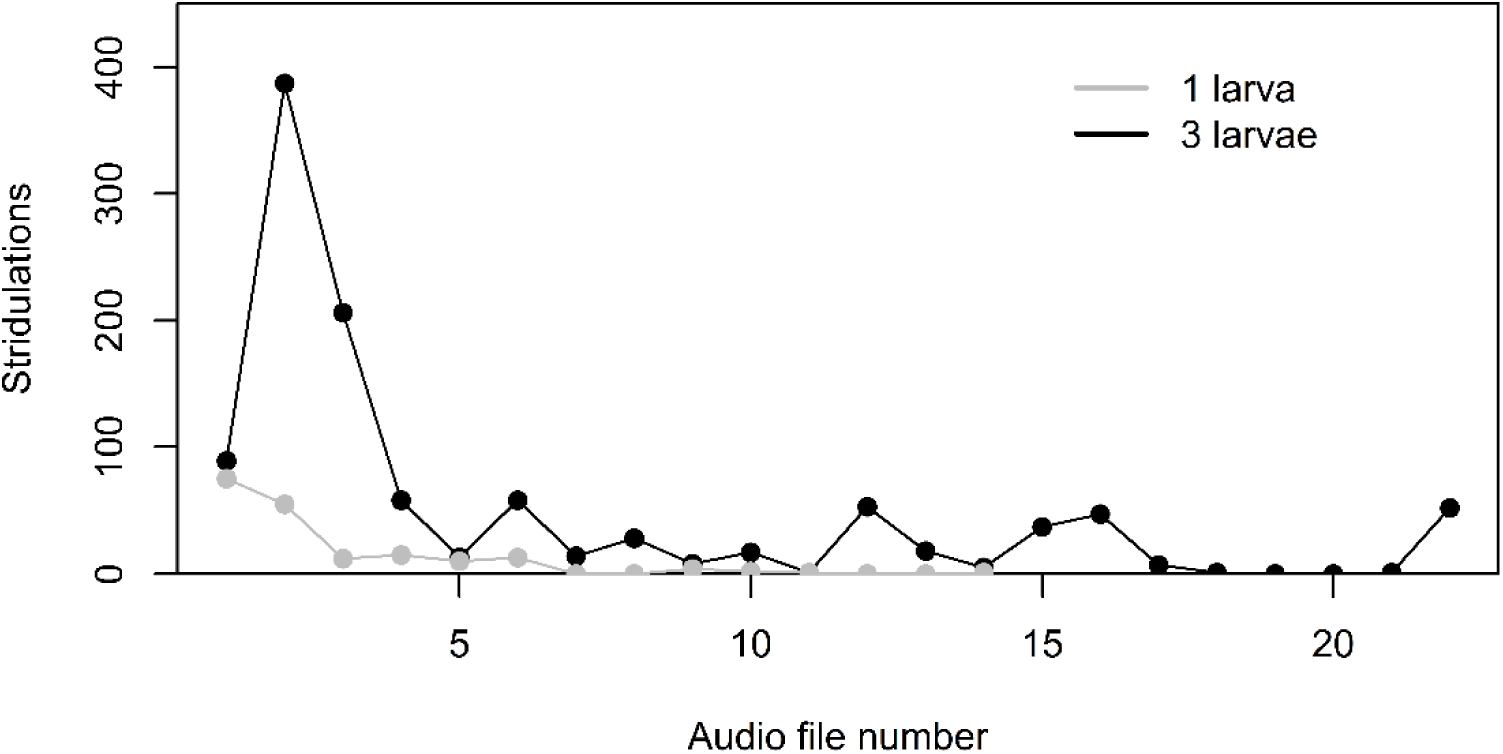
Stridulations manually counted in continuous audio recordings of third instar *Melolontha hippocastani* activities in laboratory soil incubations with 1 and 3 larvae, respectively. Each audio file was 50 min long. For the incubation with 1 larva, only 14 audio files were sequentially recorded

### Performance of data analysis routine

For the fractal dimension analysis, the chosen *f* sizes of 88.2 and 176.4 samples were best suited for describing the geometric complexity of stridulation events in the time domain, and thus for detecting them in the amplitude timeseries. For stridulation events, *FD* became more negative in comparison to incidental sounds (movement and feeding sounds, background noise, and interferences). Summing up the two *FD* time series for each analysed *sc* enhanced that effect, separating stridulations from incidental sounds even further along the y-axis. A threshold value of −4.0 was determined to be most suitable for separating stridulations from incidental sounds. Positive S*FD* values were associated with background noise and could be completely disregarded in any further analysis (Fig. 1b).

The S*FD* threshold value of −4.0 filtered out most of the incidental sounds, but for some of them S*FD* also fell below −4.0. However, the short bursts of sounds which made up one stridulation event translated into a S*FD* timeseries in which a cluster of peaks with individual peak widths of 1 or 2 *f* and spacing between peaks ranging from 2 to 10 *f* passed the threshold (Fig. 1b and 1c). For larval movement sounds and interferences, S*FD* peaks passing the threshold were usually wider than 2 *f*, and not clustered. Thus, including the second and third filter step in the data analysis routine significantly improved its capability for detecting stridulations while ignoring the majority of other sounds regardless of their origin.

The data analysis routine based on fractal dimension did not provide a direct count of stridulation events or of sound bursts within single stridulations in an audio recording. The combined 1288 stridulations in the second experiment were spread only over 893 *sc* (= 30 min) of the entire audio recordings. Stridulations were directly identified by the data analysis routine in 379 *sc* (42 %). 274 *sc* (31 %) with stridulations contained only one *SFD* peak passing the −4.0 threshold, no *SFD* peak clusters, and 240 *sc* (27 %) with stridulations were not detected. However, there was a high degree of correlation between the automatically calculated *STRAC* and the manually counted stridulations (Fig. 5).

**Fig. 5.**
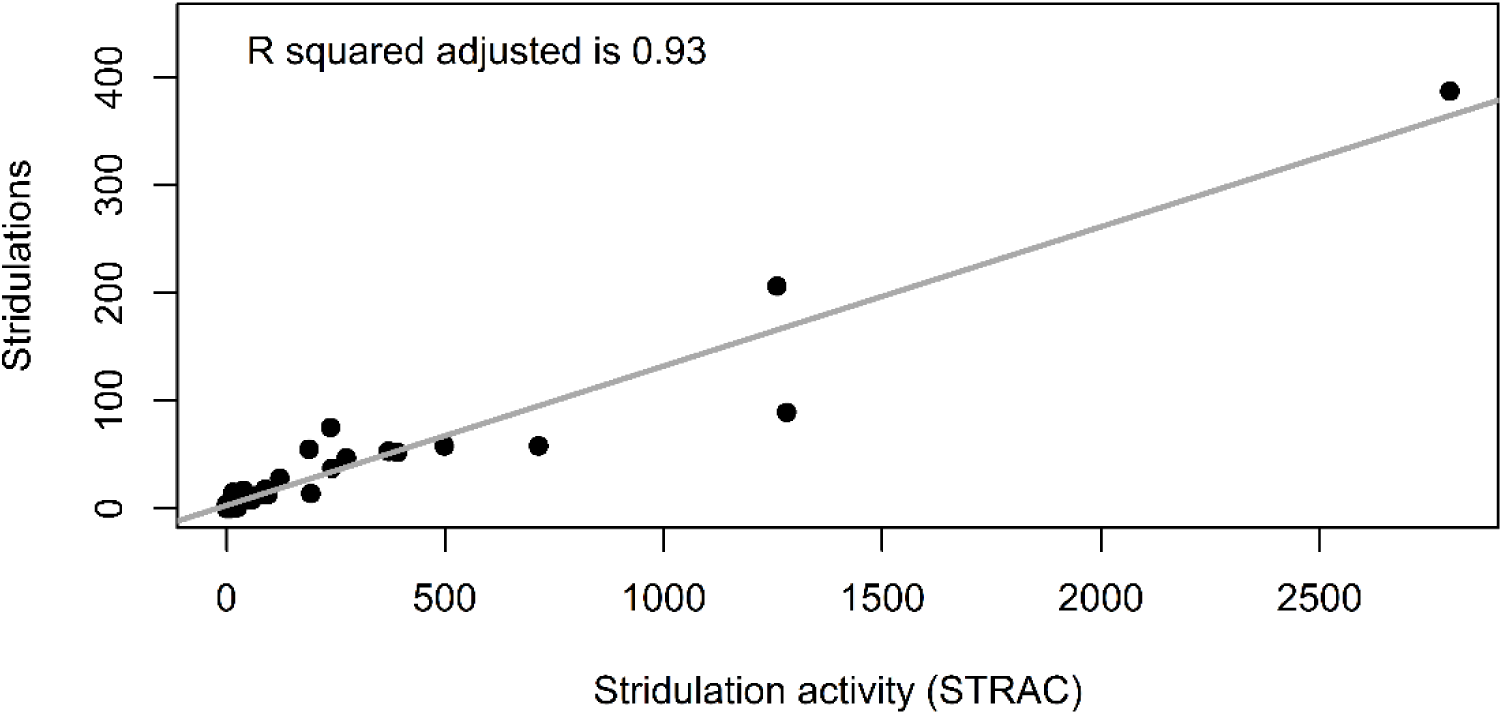
Linear regression of manually counted stridulations on automatically calculated stridulation activity (*STRAC*). Each data point came from a 50 min audio recording. The stridulation data were the same as in Fig. 4, but without differentiation between the numbers of larvae in the soil. Stridulation activity was calculated by multiplying *TI* 1 to 10 (see Fig. 1) with their respective frequencies in each 50 min audio recording and summing up the resulting products

## Discussion

It has long been known that Melolonthinae and other Scarabaeidae larvae possess stridulatory organs and morphological descriptions are available for many species (Wessel 2006). The first description for *M. melolontha* larvae stems from 1874 (Schiödte 1874). The larval stridulatory organs in Melolonthinae are located maxilla-mandibular and consist of a pars stridens (an area with fine parallel ribs) and a plectrum (sharply confined ridge) which are moving against each other (Schiödte 1874; Wessel 2006). However, actual audio recordings of larval stridulations exist only for very few Scarabaeidae species (Sprecher-Uebersax and Durrer 1998; Mankin et al. 2009b; Harvey et al. 2011). Here, we present to our knowledge the first verified audio recordings of larval stridulations from *M. melolontha* and *M. hippocastani*. The stridulations of the two species sounded quite similar, but were still distinguishable for a trained listener. The main difference was the overall duration of the single stridulations. This might have simply been the result of size differences since third instar *M. melolontha* are significantly larger than third instar *M. hippocastani* which should also result in larger stridulatory organs and possibly longer scraping times. In that case, stridulating second instar *M. melolontha* might not be distinguishable from third instar *M. hippocastani* in areas where both species co-occur. However, for pest control purposes it is already of great value to have an overview of the distribution of the genus *Melolontha* in the soil. We have no stridulation recordings of soil-dwelling Scarabaeidae larvae co-occurring with *Melolontha* spp. yet, but it has already been shown for saproxylic Scarabaeidae larvae that stridulations can be used for non-invasive species-specific monitoring (Harvey et al. 2011).

Due to the general lack of studies on beetle larval stridulations, their ecological meaning is not well understood although they are mostly interpreted as a territorial defence technique, i.e. stridulating larvae signal their presence to other larvae to avoid competition for resources and to forgo cannibalism (Kocarek 2009). Cannibalism is not uncommon (Victorsson and Wikars 1996; Kocarek 2009) and is also known for *Melolontha* spp larvae. Sprecher-Uebersax and Durrer (1998) observed in a laboratory experiment that *Lucanus cervus* larvae stridulated much more frequently directly after they were placed in a terrarium than later on which is in line with our own observations. A likely explanation is that larvae use stridulations to orient themselves in a new environment, but stridulations diminish once the larvae have settled in their new position (Sprecher-Uebersax and Durrer 1998). Apart from this, we do not really know if there are specific times in a Scarabaeidae larval life cycle during which stridulations occur more frequently than in others, if the larvae have a diurnal or seasonal rhythm, or what their complete repertoire of sounds is. In our fast screening of larvae, we only detected very few stridulations, and thus far, only one stridulation of a *M. hippocastani* larva has been recorded in the field in undisturbed soil during a survey measurement (data not shown).

A lack of detailed knowledge on the ecology of Melolonthinae and other Scarabaeidae larvae due to their cryptic nature belowground is a general problem for efficient and environmentally friendly pest control. Filling knowledge gaps on larvae population dynamics, interactions with abiotic factors, host plant preferences, attractive trap crops as well as naturally occurring pathogens has been identified as one major direction for future research to design more efficient pest management strategies (Frew et al. 2016). Such research would benefit tremendously from the timely development of non-invasive techniques for studying Scarabaeidae larvae in the field, especially of techniques facilitating real-time continuous monitoring. To date, mainly two non-invasive techniques are available for monitoring root-feeding insects in soils – X-ray microtomography and acoustic detection – of which only the latter one is directly employable in the field thus far (Johnson et al. 2007).

Since the 1900s, acoustic methods have been successfully applied in a number of insect management applications (Mankin et al. 2011; Mankin 2012). However, the focus has been mainly on airborne sounds or vibrational signals transmitted through plant parts. According to Mankin et al. (2011), only 10 root-infesting Scarabaeidae species had been acoustically studied thus far. Soil is a more challenging medium for acoustic studies in comparison to air and plant parts due to its heterogeneity. It attenuates sound more strongly, especially at high frequencies, and sound transmission can be affected by already slight changes in soil composition (e.g. bulk density, organic matter content, soil moisture, root and stone distribution) on a scale of a few centimetres (Mankin et al. 2000; Zhang et al. 2003; Mankin et al. 2011). A major challenge is still the correct identification of sounds and the differentiation between pest and nonpest signals (Mankin et al. 2000; Mankin and Lapointe 2003; Zhang et al. 2003; Mankin et al. 2009b;). However, acoustic soil studies on Scarabaeidae larvae have only focused on incidental insect sounds (feeding, movement) which were classified by comparing their spectral profiles with reference spectral profiles (e.g. Mankin et al. 2007). We present the first data analysis routine specifically targeting Melolonthinae larval stridulations in soil. It is based on the work of Schofield (2011) who was the first to use fractal dimension analysis for the detection of larval activity sounds, specifically larval feeding bites.

Fractal dimension analysis focuses on the time-domain of an audio recording, which considerably reduces its computational costs in comparison to the analysis of spectrograms and facilitates real-time acoustic identification. In addition, fractal dimension analysis is amplitude independent and thus suitable for environments with low signal-to-noise ratios like soils (Schofield and Chesmore 2008; Schofield 2011). We further developed Schofield’s approach by combining different *f* sizes, changing the *FD* calculation, and adding a new horizontal filter component. For larval feeding bites, the *f* size should ideally be equal to the length of the targeted event (Schofield and Chesmore 2008). This is not feasible for *Melolontha* spp. stridulation events which are significantly longer than feeding bites and the likelihood to falsely detect background noise increases with increasing *f* size. Instead, *f* sizes were kept small to target the single pulses within stridulations. Stridulation duration also affects the *FD* calculation. Schofield (2011) used a classical outlier detection approach by measuring fractal dimensions within a recording as distances from the mean value in multiples of the standard deviation. Stridulations take up a much larger proportion in a recording than feeding bites, significantly affecting the standard deviation around the mean. For the implementation of a vertical stridulation detection threshold, it proved more successful a) to use the median deviation around the median as a more robust measure of dispersion, and b) to separate the *FD* of stridulations and background noise further along the y-axis by combining the results of two frame sizes similarly able to capture stridulation pulses. The vertical S*FD* detection threshold separates stridulations pulses and incidental sounds with similar geometrical complexity from any other sounds. Stridulations within this subset can then be targeted by the newly developed horizontal filter, which takes into account the distinct temporal pattern of *Melolontha* spp. stridulations.

The performance of the fractal dimension analysis ultimately depends on what other sounds are present in the targeted audio recording even when applying a robust measure of dispersion as filter. The geometrical complexity of incidental sounds can vary widely and overlap with the geometrical complexity of stridulations leading to false positives, whereas stridulations can be distorted during transmission through soil in a way that they are not detectable anymore with the chosen filters. As a result, the analysis routine presented here cannot provide an absolute count of stridulation activity. However, for the laboratory experiment the result of the horizontal filter could be used to calculate a stridulation estimator (*STRAC*), which correlated very well with the manually counted stridulations. Furthermore, the number of stridulations clearly increased with increasing larval abundance. If such a relationship between the newly developed stridulation estimator and larval abundances can be verified in the field, it would allow non-invasive species-specific larval abundance measurements with a single acoustic sensor per monitoring plot for the first time. Previous studies using a single acoustic sensor to monitor incidental larval sounds were able to determine with high accuracy the presence or absence of Scarabaeidae infestations, but found only a weak correlation between sound rate and larval abundance (Mankin et al. 2001; Mankin et al. 2007). Zhang et al. (2003) managed to predict larval abundances based on incidental sounds by using four sensors at a recording point, but this set-up was more time-consuming to operate than a single sensor system.

Frew et al. (2016) advocated that for the development of improved pest management strategies for soil-dwelling Scarabaeidae larvae, it is of immediate importance to fill knowledge gaps in the ecology of these insects. Scarabaeidae larval stridulations have been neglected in soil research and our knowledge on their ecological meaning is very limited. We used *M. melolontha* and *M. hippocastani* as model organisms in the laboratory to develop the first acoustic data analysis routine specifically targeting stridulations. The fractal dimension-based method is a fast and non-compute-intensive method for pinpointing audio sections in continuous recordings in which stridulation events took place, significantly reducing the dataset for any potential further manual or automated stridulation analysis. Furthermore, it can be adjusted for detecting other acoustic events if needed by adjusting *f* sizes and the thresholds for the vertical and horizontal *SFD* filter. Acoustics should be more considered in studies of cryptic soil insects as the application of sensors in the field is simple, relatively cheap, and can provide non-destructive, continuous automated monitoring. Acoustic monitoring should not be restricted to incidental sounds, but also include stridulations to make use of its full potential for gaining significant new insights into insect ecology and biodiversity in general, and pest monitoring in particular.

## Author Contribution Statement

CMG and DC designed the experiment and analysed the data. DC designed and built the acoustic sensors. CMG performed the experiment, collected all data and developed the R script. CMG wrote the first manuscript draft and DC reviewed the manuscript.

## Ethical approval

All applicable international, national, and/or institutional guidelines for the care and use of animals were followed.

## Acknowledgements

This project has received funding from the European Union’s Horizon 2020 research and innovation programme under the Marie Sklodowska-Curie grant agreement No 703107. We would also like to thank the following persons for access to the collection sites, help during the excavation of larvae, and ecological insights into these animals: Rainer Hurling and Sabine Weldner (Northwest German Forest Research Institute).

