## Appendix S1 - R code for "Stridulations of *Melolontha spp.* larvae open up new possibilities for species-specific pest monitoring in soils"

#### Appendix S1 – R code for *Melolontha* spp. stridulation detection using fractal dimension analysis

**Author**: Carolyn-Monika Görres
**Date**: 28 April 2018

This code has been developed for the (semi-)automated analysis of the soil acoustic data recorded in the project CH4ScarabDetect. It focuses on stridulations of the species *Melolontha*, but can theoretically be adapted to detect stridulations or incidental sounds (feeding and movement sounds) of other Scarabaeidae species by simply adjusting the frame sizes in the fractal dimension calculation.

Input files for this script were pre-processed using the software Audacity (bandpass filter 200 – 5000 Hz, amplitude normalization). A user-specified number of audio files can be analysed in batch mode. Audio files in the batch will be analysed sequentially. No further actions by the user are necessary as soon as the script is running. The script can also be used if the user wants to analyse only a single audio file.

R code is highlighted in blue. Comments to the R code are highlighted in green.

Comments as well as suggestions for improving the R code are always welcome and can be send to.

##### 1. Required packages and input parameters

###### *1.1 Packages*

```
library(tuneR) # Analysis of music and speech (Ligges et al. 2016)
library(fractaldim) # Estimation of fractal dimensions (Sevcikova et al. 2014)
```

###### *1.2 Input parameters*

These settings have to be checked and if necessary changed before the script can be executed.

```
files_lowfreq = "D:/R projects/test files/" # folder containing the audio file(s) to be analysed sequentially
folder_pictures = "D:/R projects/output" # folder for the script output
duration = 2.0 # The detection of acoustic signals will be performed on time slices of a specified duration, in this case 2.0 seconds.
start_times = seq(0, 8, by = duration) # This determines the number of slices. It specifies the time (in seconds) at which the first slice should start, the time at which the last slice should start, and the duration of each slice. Has to be adjusted depending on the total length of the audio file. In this case, 5 audio slices are specified with the first one starting at second 0 and the last one starting at second 8.
audio_slices = seq(1,5, by = 1) # This list is used for indexing the single audio slices later in the script, and is just a continuous list from 1 to X. X is the total number of audio slices. If the second number in the 'start_times' code is changed, the number of audio slices has to be adjusted as well.

# IMPORTANT: If the duration is changed, then the timeArray calculation later in the script has to be adjusted as well.
```

*1.3 Batch mode*

The following code section enables R to perform functions on a single audio file. It contains the complete audio analysis code. Once an audio file has been analysed, the analysis of the next audio file will start automatically until all audio files in the specified input folder have been analysed. No further actions by the user are necessary from here on once the script is running.

```
setwd(files_lowfreq) # Sets the working directory to the folder which contains the audio files to be analysed.

fileNames <- list.files(pattern=".wav$") # Creates a list of the audio file names.

for (fileName in fileNames) { # Starts the batch mode.

    audio_lowfreq = list() # Creates a list from which the single objects can be indexed later on.

    for (i in seq_along(start_times)) {
        audio_lowfreq[[i]] = readWave(fileName,
                                from = start_times[i],
                                to = start_times[i] + duration,
                                units = 'seconds')
    } # Imports the audio file in slices of a specified length. In this example, the detection of acoustic signals will be performed on 2 s slices. If other slice sizes are desired, the input parameters in section 1.2 have to be changed.

# For the calculaton of fractal dimensions, the audio slices have to be converted into simple numeric values. We only need the amplitude data of the audio slices for this.

    amplitude_lowfreq = list()

    for (i in seq_along(audio_slices)) {
        amplitude_lowfreq[[i]] = audio_lowfreq[[i]]@left
    } # Extraction of the amplitude values from the audio slices.

# The readWave function reads .wav files as integer types. In this example, the .wav file has a 16-bit depth, this means that the sound pressure values are mapped to integer values that can range from -2^15 to (2^15)-1. The sound array can be converted to floating point values ranging from -1 to 1 as follows:

    for (i in seq_along(audio_slices)) {
        amplitude_lowfreq[[i]] = amplitude_lowfreq[[i]] / 2^(audio_lowfreq[[i]]@bit - 1)
    }

# A time representation of the sound can be obtained by plotting the pressure values against the time axis. For this it is necessary to create an array containing the time points:

    timeArray = list()

    for (i in seq_along(audio_slices)) {
        timeArray[[i]] = (0:(88200-1)) / audio_lowfreq[[i]]@samp.rate
```

```
}

for (i in seq_along(audio_slices)) {
	timeArray[[i]] <- timeArray[[i]] * 1000 # Scales time to milliseconds.
}
```

### For the calculation of fractal dimensions, each audio slice is divided into a specific number of frames. The frame size used has a major effect on the detection ability for specific sounds. It should be about the same size as the sounds which should be detected. Cockchafer larvae sounds in the soil can be grouped into three distinct groups which differ significantly in their duration: feeding sounds, stridulation sounds, and moving sounds. The smaller the chosen frame size(s), the longer the calculation time for each slice. Thus, the selected frame sizes are a compromise. Cockchafer stridulation sounds seem to be detected best by using frame sizes of 88.2 and 176.4 (2 and 4 milliseconds, respectively.)

```
fractalDimensions_88.2 = list() # List for the fractal dimensions of the low frequency audio slices with a frame size of 88.2.
fractalDimensions_176.4 = list() # List for the fractal dimensions of the low frequency audio slices with a frame size of 176.4.
```

### Calculation of fractal dimensions for each frame size. Frames do not overlap. “Madogram” is used as estimator for the fractal dimension. According to Gneiting et al. (2010), this estimator is recommended based on robustness and efficiency.

```
for (i in seq_along(audio_slices)) {
	fractalDimensions_88.2[[i]] = fd.estimate(amplitude_lowfreq[[i]],
	                                          methods="madogram",
	                                          window.size=88.2)
}
for (i in seq_along(audio_slices)) {
	fractalDimensions_176.4[[i]] = fd.estimate(amplitude_lowfreq[[i]],
	                                           methods="madogram",
	                                           window.size=176.4)
}
```

### Sounds in the audio slice can be regarded as outliers and fractal distances are calculated as a means to detect these outliers. In contrast to Schofield (2011), this script does not rely on the mean and standard deviation, but calculates the "median deviation from median" (MAD) for each frame (Leys et al. 2013). The median works better when sounds take up a significant portion of a slice.

```
FractalDistance_88.2 = list() # List for the fractal distances of the low frequency audio slices with a frame size of 88.2.
FractalDistance_176.4 = list() # List for the fractal distances of the low frequency audio slices with a frame size of 176.4.
```

### Important: The script does not take the absolute value when subtracting the median fractal dimension from the fractal dimension of a frame.

```
for (i in seq_along(audio_slices)) {
```

```
        FractalDistance_88.2[[i]] = (fractalDimensions_88.2[[i]]$fd -
                median(fractalDimensions_88.2[[i]]$fd)) /
                mad(fractalDimensions_88.2[[i]]$fd)
}

for (i in seq_along(audio_slices)) {
        FractalDistance_176.4[[i]] = (fractalDimensions_176.4[[i]]$fd -
                                        median(fractalDimensions_176.4[[i]]$fd)) /
                                        mad(fractalDimensions_176.4[[i]]$fd)
}
```

### The fractal distances for the different frame sizes are brought together into one number for the detection of acoustic signals. In a first step, the timeseries for the different frame sizes are interpolated to the same time interval (2 ms). In a second step, the sum of the timeseries is calculated. The fractal distances timeseries with a frame size of 88.2 consist of 1000 samples each. The results of the other frame size is interpolated through time to reach the same amount of samples.

```
Interpol_176.4_x <- seq(1,1000, by=2) # Time sequence for the subsequent linear interpolation.
Interpol_FD176.4 = list() # List for the interpolations of the fractal distances with frame size 176.4.

for (i in seq_along(audio_slices)) {
        Interpol_FD176.4[[i]] <- approx(Interpol_176.4_x, FractalDistance_176.4[[i]],
                                        n=1000)
        Interpol_FD176.4[[i]] <- Interpol_FD176.4[[i]]$y
}

FractalDistance_Sum = list()

for (i in seq_along(audio_slices)){
        FractalDistance_Sum[[i]] <- (Interpol_FD176.4[[i]] +
                                        FractalDistance_88.2[[i]])
} # Summing up the two types of fractal distances.
```

### Using fractal distance alone, it is possible to detect larval sounds in an audio file, but it is not possible to distinguish between stridulation and moving sounds. However, stridulation sounds consist of evenly spaced pulses whereas moving sounds have no recognizable, repeating pattern. For the detection of stridulations in the fractal distances, the following R code determines the starting point of each event passing a threshold of -4, and then calculates the distance between adjacent detected events. In contrast to movement sounds, stridulation sounds contain several events (= pulses) which are less than 10 samples apart.

```
FD_rle <- lapply(FractalDistance_Sum, function(i){rle(as.numeric(i) <= -4.0)})
# rle (run length encoding) computes the lengths and values of runs of equal values in a vector. Look for runs containing values <= -4.0

myruns = list()

for (i in seq_along(audio_slices)){
```

```
		myruns[[i]] <- which(FD_rle[[i]]$values == TRUE & FD_rle[[i]]$lengths <= 2)
	} # Finds indices of the runs with sample length <= 2.

# The following code finds the end of each detected run.

	runs.lengths.cumsum = list()

	for (i in seq_along(audio_slices)){
		runs.lengths.cumsum[[i]] = cumsum(FD_rle[[i]]$lengths)
	}

	ends = list()

	for (i in seq_along(audio_slices)){
		ends[[i]] = runs.lengths.cumsum[[i]][myruns[[i]]]
	}

	Rle_diff <- lapply(ends, function(i){c(0, diff(i))}) # Calculates the difference between
	adjacent runs. Single pulses of stridulation should be close together.

# The following code creates for each audio slice a figure showing the amplitude versus time,
the summed fractal distance versus time, and the audio signal detection result.

	setwd(folder_pictures) # Sets the working directory to the folder which should contain
	the R output.
	dir.create(fileName) # Creates a subfolder for the audio file currently being analysed.
	setwd(fileName) # Sets the working directory to this subfolder.

	for (i in seq_along(audio_slices)) {
		png(paste("Frame", audio_slices[[i]], " FracDistSum.jpg", sep = ""),
		width = 1200, height = 960)
		par(mfrow=c(3,1))
		plot(timeArray[[i]], amplitude_lowfreq[[i]], type='l', col='black'
		, xlab='Time (ms)', ylab='Amplitude (low frequency)', main=audio_slices[[i]])
		plot(FractalDistance_Sum[[i]], type='n', ylim=c(-6,2), main="Fractal distance –
		sum of frame sizes 88.2 and 176.4")
		lines(FractalDistance_Sum[[i]])
		abline(-4.0,0.0, col="blue", lwd=2)
		tryCatch({plot(ends[[i]],Rle_diff[[i]], ylim=c(0,10), pch=19, cex=3.0, col='red',
		xlim=c(0,1000))},
		error=function(e) plot(0,type='n',axes=FALSE,ann=FALSE)) # This plot is not
		plotted when no events were detected.
		abline(6.0,0.0, col="blue")
		par(mfrow=c(1,1))
		dev.off()
	}

# The next figure summarizes the results for the Rle_diff calculation, i.e. counts the number of
adjacent runs which were less than 10 samples apart. For this it is necessary to first determine
which elements in list "Rle_diff" have more than one element. These are the frames with peaks
detected using the Rle_diff criteria.
```

```
Ends_length = list()

for (i in seq_along(audio_slices)) {
        Ends_length[[i]] <- length(ends[[i]])
}

Frames <- which(Ends_length != 0)
Frames <- length(Frames) # Number of frames with peaks detected using the Rle_diff criteria.

# Counts the number of events for each value of Rle_diff between 1 and 10.

Rle_diff2 <- unlist(Rle_diff)
count1 <- length(which(Rle_diff2 == 1)) # Counts the number of events for a distance of 1 sample.
count2 <- length(which(Rle_diff2 == 2))
count3 <- length(which(Rle_diff2 == 3))
count4 <- length(which(Rle_diff2 == 4))
count5 <- length(which(Rle_diff2 == 5))
count6 <- length(which(Rle_diff2 == 6))
count7 <- length(which(Rle_diff2 == 7))
count8 <- length(which(Rle_diff2 == 8))
count9 <- length(which(Rle_diff2 == 9))
count10 <- length(which(Rle_diff2 == 10))

Bottom_axis <- c(0, 1,2,3,4,5,6,7,8,9,10) # Numbers for the x-axis in the following figure.
Count_axis <- c(Frames, count1, count2, count3, count4, count5, count6, count7, count8, count9, count10) # Creates the y-data for the following figure.

png(paste("Result of Rle_diff calculation.jpg", sep = ""), width = 1200, height = 960)
plot(Bottom_axis, Count_axis, type='l', xlab='Distance between adjacent runs (# of samples)', ylab='Count of events occuring', col='blue', lwd=2, main='Result of Rle_diff calculation')
dev.off()

Counts <- cbind(Bottom_axis,Count_axis) # Presents the count results as data table.
write.csv(Counts, "Count result.csv", row.names=FALSE) # Exports the data table as .csv file

setwd(files_lowfreq) # Set the working directory again to the folder which contains the audio files to start the analysis of the next audio file.

} # Stops the batch mode once all audio files have been analysed. A warning will appear which can be ignored.
```

#### 2. Additional information

##### *2.1 Additional information*

This R code was executed with the following settings:

```
## R version 3.4.3 (2017-11-30)
## Platform: x86_64-w64-mingw32/x64 (64-bit)
## Running under: Windows 10 x64 (build 17134)
##
## Matrix products: default
##
## attached base packages:
## [1] stats graphics grDevices utils datasets methods base
##
## other attached packages:
## [1] fractaldim_0.8-4 abind_1.4-5 tuneR_1.3.2
##
## loaded via a namespace (and not attached):
## [1] Rcpp_0.12.14 digest_0.6.15 rprojroot_1.3-1 MASS_7.3-47
## [5] backports_1.1.2 signal_0.7-6 magrittr_1.5 evaluate_0.10.1
## [9] stringi_1.1.6 rmarkdown_1.8 tools_3.4.3 stringr_1.2.0
## [13] yaml_2.1.16 compiler_3.4.3 htmltools_0.3.6 knitr_1.17
```

##### *2.2 References*


Gneiting T, Sevcikova H, Percival, DB (2010) Estimators of fractal dimensions: Assessing the roughness of time series and spatial data. University of Washington, Department of Statistics, Technical Report No. 577, pp. 38.

Leys C, Ley C, Klein O, Bernard P, Licata L (2013) Detecting outliers: Do not use standard deviation around the mean, use absolute deviation around the median. J Exp Soc Psychol 49:764–766. https://doi.org/10.1016/j.jesp.2013.03.013

Ligges U, Krey S, Mersmann O, Schnackenberg S (2016) tuneR: Analysis of music. http://r-forge.r-project.org/projects/tuner/. Accessed 21 August 2018.

Schofield J (2011) Real-time acoustic identification of invasive wood-boring beetles. Dissertation, University of York, Department of Electronics, York, UK, pp. 176.

Sevcikova H, Percival D, Gneiting T (2014) fractaldim: Estimation of fractal dimensions. R package version 0.8-4. https://CRAN.R-project.org/package=fractaldim. Accessed 21 August 2018.

**Example output figure 1** Detection of cockchafer larvae stridulations using fractal dimension analysis (see R code for details). Fig. 1a: Audio recording with two stridulations (at ~650 ms and ~1100 ms) and larval moving sounds (from ~1700 ms onwards). Fig. 1b: Summed fractal distance (*SFD*) for every 2 ms (=frame) of the audio recording. Peaks crossing a threshold of -4.0 are first indicators of stridulation events. Fig. 1c: Number of frames between adjacent peaks (*TI*) crossing the threshold in Fig. 1b. A distance of less than 10 frames is indicative of a peak of clusters crossing the threshold in Fig. 1b, and thus a stridulation event

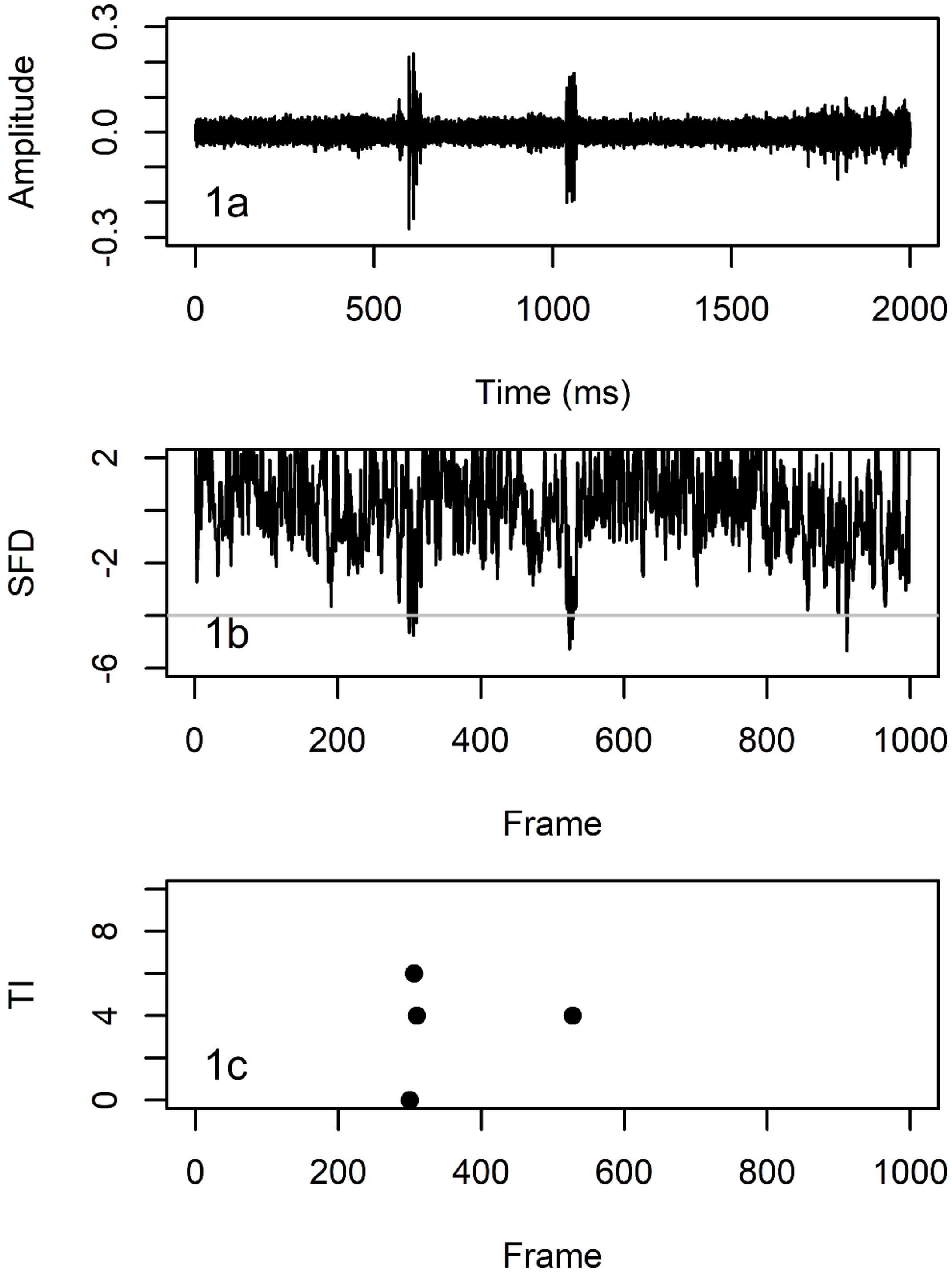

**Example output figure 2** Example of count results

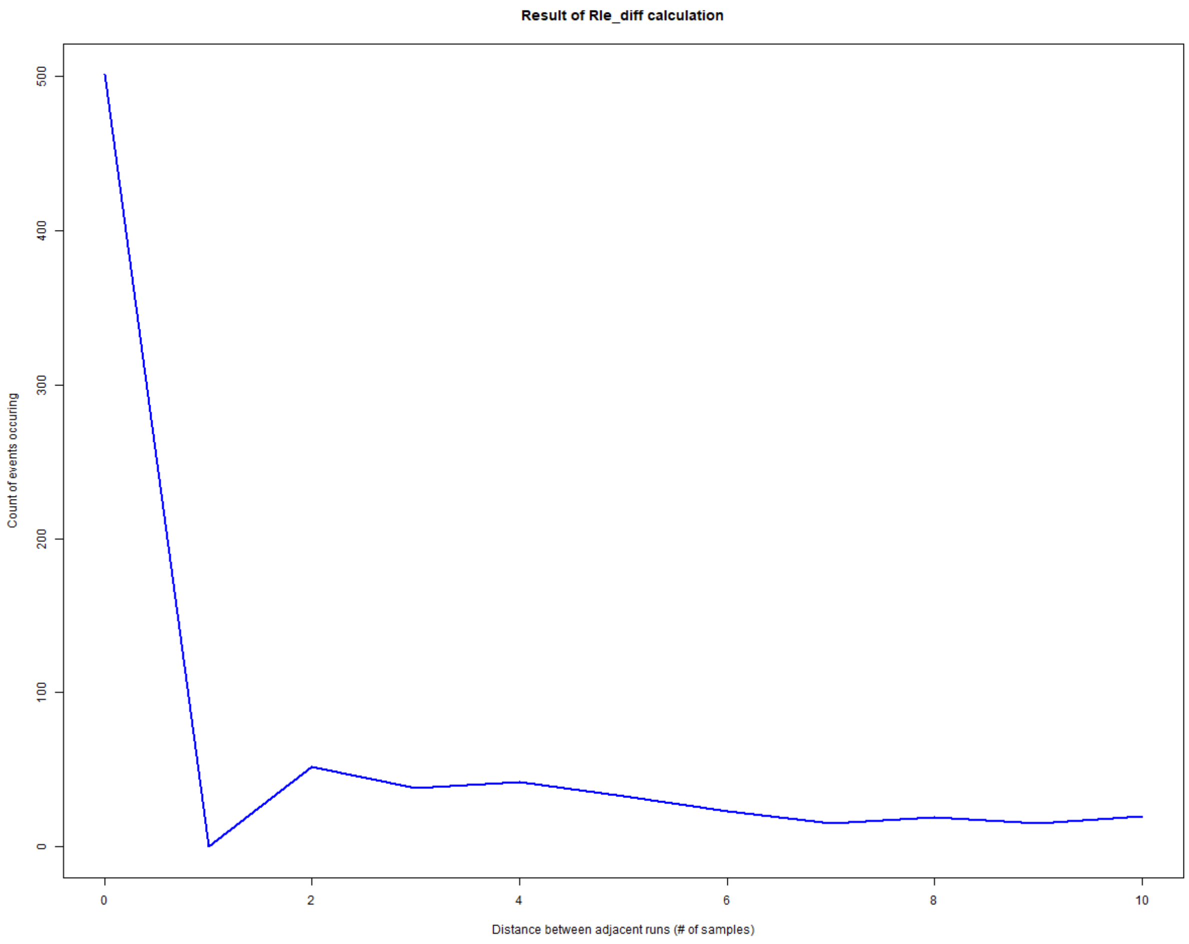
